# Deep learning of Protein Sequence Design of Protein-protein Interactions

**DOI:** 10.1101/2022.01.28.478262

**Authors:** Raulia Syrlybaeva, Eva-Maria Strauch

## Abstract

**Motivation:** As more data of experimentally determined protein structures is becoming available, data-driven models to describe protein sequence-structure relationship become more feasible. Within this space, the amino acid sequence design of protein-protein interactions has still been a rather challenging sub-problem with very low success rates - yet it is central for the most biological processes.

**Results:** We developed an attention-based deep learning model inspired by algorithms used for image-caption assignments for sequence design of peptides or protein fragments. These interaction fragments are derived from and represent core parts of protein-protein interfaces. Our trained model allows the one-sided design of a given protein fragment which can be applicable for the redesign of protein-interfaces or the *de novo* design of new interactions fragments. Here we demonstrate its potential by recapitulating naturally occurring protein-protein interactions including antibody-antigen complexes. The designed interfaces capture essential native interactions with high prediction accuracy and have native-like binding affinities. It further does not need precise backbone location, making it an attractive tool for working with *de novo* design of protein-protein interactions.

**Availability:** The source code of the method is available at https://github.com/strauchlab/iNNterfaceDesign

**Supplementary information:** Supplementary data are available at Bioinformatics online.

## 1 Introduction

The ability to computationally engineer protein sequences has a wide range of applications from therapeutics (Fosgerau & Hoffmann, 2015; Khera & Maity, 2019), to vaccines (Li & Li, 2020; Liu et al., 2020; Malonis et al., 2020; X. Zhou et al., 2020), sensors (Karimzadeh et al., 2018; Merkx et al., 2019) or protein-based materials (Capezza et al., 2019; de la Rica & Matsui, 2010) and beyond that. While there has been progress towards designing protein folds, much improvement is needed for the redesign or *de novo* design of protein-protein interfaces (PPIs). The success rates for the *de novo* generation of protein-protein interactions archived by existing methods is very low, with only a few examples demonstrating it is possible (Cao et al., 2020; Fleishman et al., 2011; Strauch et al., 2014). Even most recent work illustrates an average success rate of 0.25% while requiring substantial computational and laboratory resources (Cao et al., 2021) underlining that it is still highly challenging. With DeepMind using deep learning algorithms and entering the field of protein structure prediction, much has changed about how machine learning approaches can be used for protein folding and also design. Recent work using neural networks improved substantially accuracy in structure prediction (Baek et al., 2021; Jumper et al., 2021; Senior et al., 2020) and protein sequence design (Anand et al., 2020; Chen et al., 2020; Gao et al., 2020; O’Connell et al., 2018). The latter methods outperform traditional methods for sequence design based on energy function integrated into sampling, filtering, and optimization procedures (Adolf-Bryfogle et al., 2018; Desjarlais & Handel, 1995; Raha et al., 2000). The average sequence recovery achieved by the current top-performing protein-design programs is around 30% (J. Zhou et al., 2020), while SPROF model (Chen et al., 2020) achieved 39.8% on independent test sets. Based on these inspiring results, we developed a deep learning-based approach for the sequence design of PPIs which is a sub-problem within the sequence design space.

A main challenge for design of new protein interfaces is the identification and placement of energetic “hot-spots” (Cukuroglu et al., 2014; Wells & Clackson, 1995). A characteristic feature of protein interactions is that interface residues do not contribute equally to the binding energy and only a few residues have energetically highly favorable interactions. Their placement is complicated by the diversity a given rotamer can have or the conformational states a protein can exhibit. Further, models should consider both surface contacts with the target protein, but also protein stability of the unbound state. Here, we report on the recovery of small patches of PPIs containing hot-spot residues. We focus on design of amino acid sequences (AAS) of “sixmer” (six residue long) fragments of protein ligands exhibiting high affinity PPIs. Our method is intended to be used for the recovery of native AAS for a fixed backbone of protein ligand or the design of *de novo* interactions. For the latter case, often, the precise location of the backbone of the ligand is not exactly known which may also be applicable for the re-design of complexes that have been generated through homology modeling. In these cases, deviations in bond lengths, valent or dihedral angles may occur and need to be addressed. To mimic this challenge, all ligands were perturbed before the evaluation of our sequence design model.

Our method differs from existing deep learning models for design of PPIs (Wu et al., 2021; Zhang et al., 2021), which are intended to help docking by defining interaction pairs of residues of two counterparts rather than creating the counterparts itself. The architecture of neural network is inspired and based on a model generating captions of an image with visual attention (Xu et al., 2015) turning pictures into captions. Hence, we treat the structures of protein as the 3D object to be captured. Features are then extracted using machine learning vision techniques and then translate these into sequences specifically focusing on the properties of how its counterpart (protein ligand) should look like. We developed two versions of the deep learning models, named PepSeP1 and PepSeP6, producing single and multiple outputs correspondingly. The methods significantly outperform Rosetta’s FastDesign mover (Khatib et al., 2011; Tyka et al., 2011) on an independent test set. Comparison of binding affinities of native and designed peptides has shown small difference of 0.7 Rosetta energy unit (REU) in favor of native structures according to estimation by means of the current Rosetta scoring function (ref2015,(Alford et al., 2017). As mentioned, the developed model is intended to be applied on difficult cases involving backbones even far from native geometries for which most current methods are not very adapted for. An extension of the method allows the design of larger fragments by making subsequent designs of connected backbone fragments. Considering these results, we provide a highly promising neural network (NN)-based approach for the design of PPIs.

## 2 Materials and methods

### 2.1 Test Sets

The method is trained and tested on peptide–binding site complexes, extracted from native PPIs (Fig. 1). The peptide of the complex is 6-residue fragment of a protein ligand and the binding site is a patch of a receptor consisting of 24-48 residues which are closest to backbone atoms of the 6-residue fragment. All complexes for this study originate from multichain structures obtained from Protein Data Bank(Berman, 2000).

**Figure 1.**
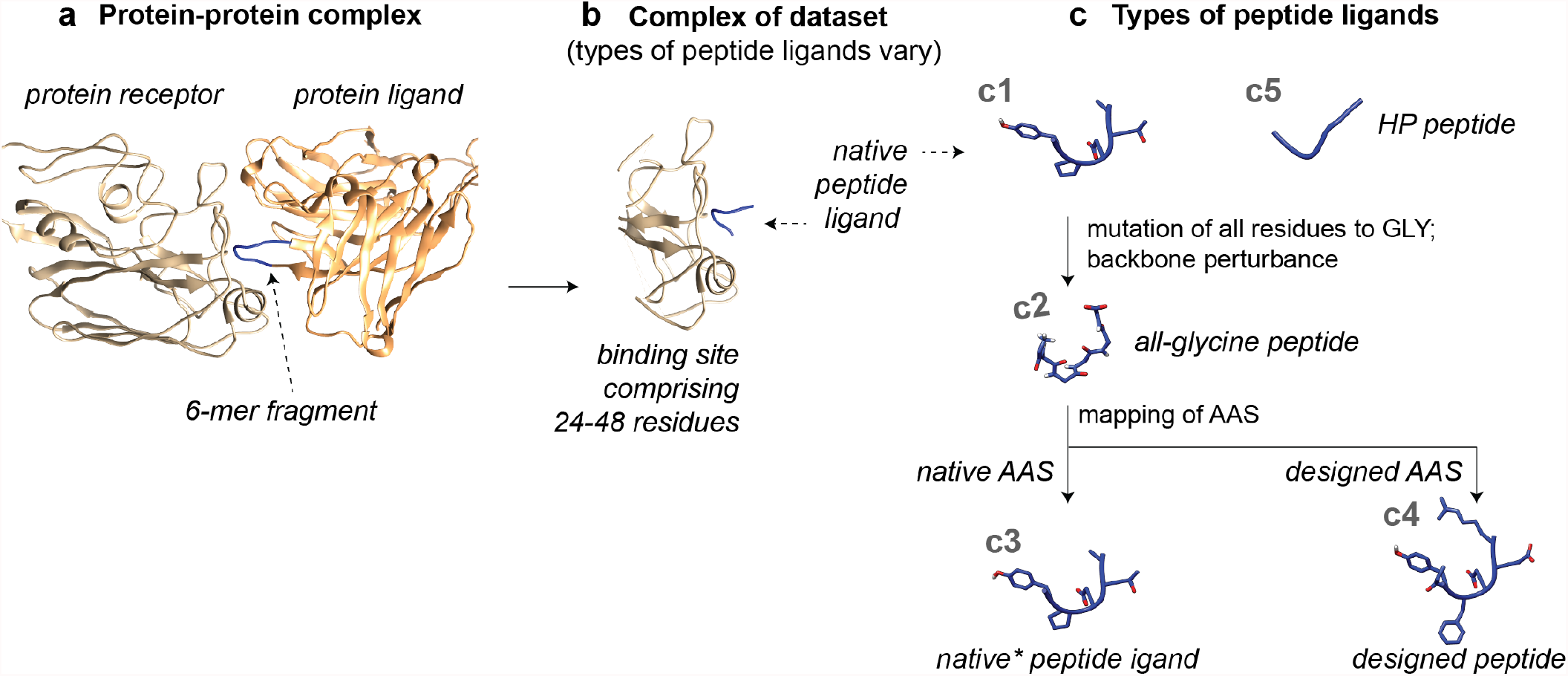
Generation of datasets of peptide–binding site complexes and overview of structures utilized in the study. **(a)** Crystal structure of protein-protein complex with selected interacting 6-residue fragment of protein ligand. **(b)** Peptide–binding site complex of datasets. **(c)** Types of peptide ligands and their generation: native peptide ligand which is the 6-residue fragment depicted in the subfigure **a** (c1), was mutated into an all-glycine peptide ligand and perturbed (c2, SI 1.2). Perturbed backbones were then either reverted to their native amino acid sequence (c3, and annotated as native* within main text to highlight the backbone perturbation) or designed with PepSeP1 (c4). HP (highly perturbed) backbones (c5) have non-native backbone geometry after extensive perturbation. s

The complexes were extracted from 9,101 PDB-files. 70 Files containing complexes of influenza’s hemagglutinin (HA) and coronaviruses MERS-CoV, SARS-CoV and SARS-CoV-2 proteins with antibodies (Table S2) were separated from the main set to form a benchmark set, all other complexes with the listed antigens were deleted from the main set. The files of the main set were randomly split to construct training, validation and test sets in quantities of 8 581, 271, 179 respectively; total number of complexes obtained from these crystal structures are 95 087, 2 933 and 1 261; part of the curation was to eliminate any duplicates of the complexes to enable independent training and test sets. The benchmark set, omitting duplicates, consists of 915 complexes which are independent of the main test set regarding antigens; it contains antibody or antibody fragments which can be present in other datasets.

Our method was evaluated using both test and benchmark sets, referred as “set T” and “set B” respectively. Set T is divided into two subsets T*-*ho and T*-*he, based on whether they were derived from homo- and hetero-oligomeric PPIs correspondingly. Set B was used for evaluation of sequence recovery of interfaces focusing on antibody-antigen protein complexes. This set is divided into subsets as well (Table 1). Quantity of complexes in the benchmark subsets depending on types of antigens is summarized in Table S3. Other details related to construction of the datasets can be found in methods section of SI (SI 1.1 – 1.3).

**Table 1.** Subsets of the benchmark set

| Subset | Source of a backbone fragment (peptide) to be designed | Source of a binding site to which the peptide is attached | Number of complexes |
| --- | --- | --- | --- |
| B-ab/ag | Antibody | Antigen | 150 |
| B-ag/ab | Antigen | Antibody | 144 |
| B-ab/ab | Antibody | Antibody | 485 |
| B-ag/ag | Antigen | Antigen | 136 |

#### Generation of highly perturbed (HP) peptide fragments

Beside testing the methods on all-glycine peptide ligands extracted from native interactions, we tested our method on artificially highly perturbed (HP) peptide variations of a 6-residue all-glycine peptide ligands fragment (Fig. S3). These backbone variations were generated by PepBB deep learning model, developed by our group and soon to be published (available at (*INNterfaceDesign*, 2021)), as a starting point of the complexes of the test set T. Twelve peptide ligands were generated for each of the binding sites, the ones with the lowest RMSD to native peptide ligands were selected as HP peptides. Average RMSD of them is about 5.0 Å (3.8 Å for non-terminal residues). High RMS difference are due to 20% of HP peptides with a reverse orientation to the native conformation: they are on same location but N- and C- termini are other way around (Figure S3a). Terminal residues of the native peptide ligands are typically non-interacting (SI 1.2) which also contributed to RMSD. Due to some deviations from native backbones in terms of subtle geometrical features, they provide very challenging targets for AAS design and recovery of hot-spot interactions.

### 2.2 Input data for deep learning models

Input for the neural networks are geometrical and sequence features of the binding sites and XYZ coordinates of backbone atoms of the peptide ligands. We utilized two versions of the binding sites: first is based on all initially selected residues (24-48), and a second reduced representation, consisting of 20 residues of the binding site closest to the peptide ligands for improved accuracy. As a geometrical feature description of the structures, we used distance maps with either N or O backbone atoms. For each of these two atom types, we generated an intra- (map 1) and intermolecular distance map (map 2), describing either distances for the interface atoms or across the interface respectively. Input arrays and the corresponding descriptors are summarized in Table 2 (details under SI 1.4). Input 5, not mentioned in Table 2, contains binary values (0 or 1) denoting the type of PPI (homo- or hetero-oligomeric).

**Table 2.** Input from structures used for feature generation for deep learning models during training and testing.

| Descriptor |  | Binding site | Peptide | Reduced binding site |
| --- | --- | --- | --- | --- |
| Topological | Distance maps 1 (intramolecular) |  | Input 4 | Input 9 |
|  | Distance maps 2 (intermolecular) | Input 1 |  | Input 7 |
|  | Secondary Structure | Input 3 |  | Input 8 |
| AAS |  | Input 2 | Input 6 |  |

### 2.3 Architecture of Developed Neural Networks

For our sequence design algorithm, we developed a two-directional architecture involving two neural networks. The main block produces AAS for 6-residue peptide backbones (output 1) based on input 1-5 (containing AAS of the binding site and topological features of the interfaces, Table 2 and SI 1.4). To improve its accuracy, we developed the second module (reverse prediction block) trained on input 5-9, which will produce output 2, a reduced representation of the target binding site. Hence, output 2 should mirror most of input 2. Training conditions required not only high rates of recovery of native AAS of the peptide, but it also required high accuracy for output 2 when using output 1 in the reverse prediction block as input, instead of input 6 (Fig. 2). The combination of these two neural networks improved sequence recovery and also resulted in more favorable binding energies (> 4 REU improvement) making our model comparable to native interactions. After training, the reverse prediction block becomes redundant, whereas the trained main block will be used for predictions of PepSeP1.

**Figure 2.**
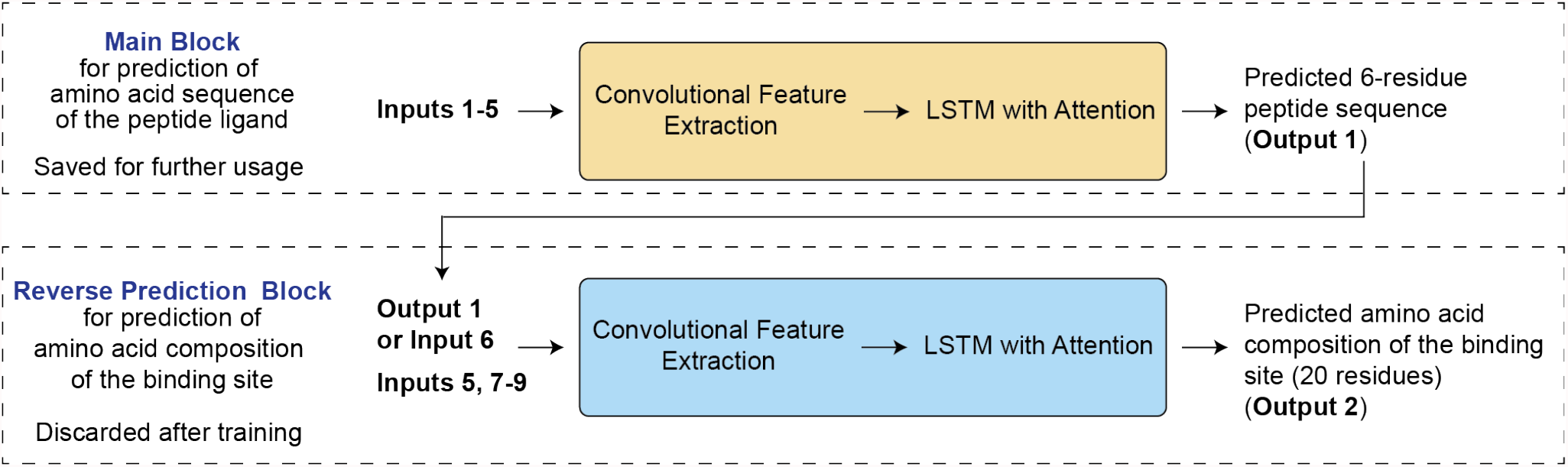
Peptide ligand sequence recovery method by PepSeP1 attention neural network. There are 9 types of input arrays provided to the neural networks (see details under 2.2 and SI 1.4).

Both blocks have an encoder-decoder architecture which has been successfully utilized for sequence prediction models (Chen et al., 2020). The architecture of the blocks is very similar and loosely based on an TensorFlow tutorial (Image Captioning with Visual Attention, n.d.) about how to implement models generating image captions with visual attention (Xu et al., 2015), specifically the decoder part. The encoders are based on convolutional layers. The PepSeP6 variation of our algorithm, producing six predicted AAS, uses the same infrastructure as PepSeP1. The scheme of PepSeP6 is presented in Fig. S4. Five outputs are generated by passing feature vectors produced by the encoder of the main block of PepSep1 model to the decoder of PepSeP6 five times; each of five iterations is accompanied by a final hidden state from the previous prediction of the sequence. The sixth sequence is the output of the PepSep1 model, which is provided to PepSeP6. A single peptide ligand sequence predicted by PepSeP6 is randomly selected from the resulting sequences after and forwarded to the reverse prediction block of PepSeP1 providing the bidirectional approach. The neural network was built in TensorFlow using Keras API (more details under SI 1.4-1.6).

### 2.4 Assessment of experimental and predicted peptide ligands

#### Refinement and calculation of binding energies of complexes of test sets

The target binding sites were relaxed without peptide ligands. Pertubed peptides were then added back either with their native or redesigned AAS (Fig.1). Optimization of side chain conformations of all residues of the complex and subsequent refinement of poses of the peptide ligands was done applying the FastRelax mover three times over 300 step while applying harmonic constraints (standard deviation SD = 1.0) for peptide ligands based on perturbed native-like backbones (c2). Less restrictive constraints (SD=3.0 with width parameter of 2.0 Å) were used for HP peptides (c5) predictions. Binding free energies of the complexes were estimated using InterfaceAnalyzerMover (Stranges & Kuhlman, 2013) with repacking chains after separation. All stages of the refinement were carried out three times using FastRelax mover over 300 steps. The structure with the lowest score out of three results was selected for the next operation. The Rosetta scoring function ref15 was used for all calculations.

#### Redesign of complexes

Peptide ligands designed by PepSeP1 method underwent an additional design step after refinement using the FastDesign protocol to compare results with original performance of PepSeP1. We set constrains on residue types according to position-specific scoring matrices (PSSM) based on outputs of PepSeP1 (SI 1.7). We performed three different protocols of redesign. We controlled how many possible amino acid types are allowed at each position. Variations are denoted as RD3, RD5 and RD20, reflecting whether either the most probably three or five amino acid types should be used or whether all amino acid were allowed. Probabilities were applied in form of described PSSMs extracted from the PepSeP1 model. In comparison, we also performed redesign using the FastDesign protocol on all-glycine peptide backbones. Two types of relax scripts were utilized during the redesigns: default (MonomerRelax2019) and InterfaceDesign2019.

## 3 Results

Table 3 summarizes performances of the developed deep learning models alongside with results of different approaches based on FastDesign protocol(Maguire et al., 2021; Tyka et al., 2011) on recovery of native sequences of the peptide ligands of the test and benchmark sets.

**Table 3.** Rates of recovery of native residues of peptide ligands of sets T and B achieved by PepSeP1 and PepSeP6 methods based on all-glycine peptides, obtained after redesigning of PepSeP1 sequences by means of Rosetta’s FastDesign protocol according to RD3, RD5, RD20 schemes and obtained by FastDesign protocol (MonomerRelax2019)) applied to all-glycine peptide fragments.

| Subset | $R_{all}$, % | | | | | | $R_{hot-spot}$, % | | | | | |
| --- | --- | --- | --- | --- | --- | --- | --- | --- | --- | --- | --- | --- |
|  | PepSeP1 | RD3 | RD5 | RD20 | PepSeP6 | FastDesign | PepSeP1 | RD3 | RD5 | RD20 | PepSeP6 | FastDesign |
| T-ho | 51.55 | 41.87 | 36.53 | 29.73 | 45.61 | 23.35 | 57.75 | 56.23 | 55.18 | 50.41 | 51.09 | 44.18 |
| T-he | 28.56 | 27.22 | 26.1 | 22.64 | 26.02 | 24.93 | 37.19 | 41.81 | 44.01 | 40.34 | 31.7 | 46.67 |
| **T** | **44.42** | **37.34** | **33.31** | **27.54** | **39.54** | **23.84** | **51.07** | **51.58** | **51.58** | **47.16** | **44.79** | **44.98** |
| B-ab/ag | 28.11 | 22.98 | 23.99 | 23.11 | 24.7 | 22.7 | 33.52 | 31.41 | 32.05 | 33.33 | 30.72 | 31.74 |
| B-ag/ab | 18.06 | 20.95 | 21.43 | 21.43 | 16.22 | 23.24 | 24.14 | 33.63 | 30.97 | 45.13 | 19.37 | 48.67 |
| B-ag/ag | 56.25 | 42.03 | 37.87 | 28.43 | 47.75 | 22.43 | 75.52 | 69.27 | 64.58 | 55.73 | 62.44 | 40.62 |
| B-ab/ab | 65.53 | 54.7 | 47.65 | 37.61 | 61.4 | 25.14 | 77.16 | 73.89 | 70.87 | 65.01 | 72.69 | 50.43 |

Sequence recovery accuracy (*R*_*all*_) on set T is 44.42%. The results depend greatly on types of PPIs: native sequence recovery rates for homo- and hetero-oligomeric PPIs are 51.55% and 28.56%, respectively. Better performance of the method on obligate PPIs is affirmed by accuracy exceeding 56% archived on subsets B-ab/ab and B-ag/ag. Lower rates archived on hetero-oligomeric PPIs are likely due to fact that substantial part of heteromeric protein interfaces are transient interactions and thereby weaker in nature than assembled complexes. Further, residue composition of transient interactions is more diverse and include higher rates of polar and charged groups alongside hydrophobic amino acids(Acuner Ozbabacan et al., 2011). Ligand residues of antigen-antibody interfaces of subsets B-ab/ag and B-ag/ab, which are transient PPIs, are predicted with 28.11% and 18.08% success rate. Antigen structures are represented as a single chain or as a fragment of the chain in the original crystal structures often. Usage of the method on the benchmark set where all hemagglutinin complexes are extracted from trimer structures (the coronavirus complexes are same as in the main benchmark set) provide overall accuracy for subset B-ab/ag of 31.85 %. Detailed data regarding the accuracy of the method depending on types of antigens can be found in Table S4, best results for PepSeP1 are observed in case of SARS-CoV-2: 41.67% and 50% in subsets B-ab/ag and B-ag/ab, correspondingly. Secondary structure of the peptide fragment impacts the accuracy as well (Table S5): the highest recovery rate of 47.11% was observed for β-sheet structures, accuracies of predictions for a-helices and especially for loops were lower (46.17 and 39.35%, respectively).

The performance of the model on recovery of hot-spot residues of the peptides was considered as well (Table 3). Energy contributions of individual residues to the binding ΔΔ*G*_*i*_ were obtained by alanine scanning (Kortemme et al., 2004). Contacts were treated as hot-spots if they had an binding energy contribution of at least 3 REU. Success rate *R*_*hot-spot*_ was calculated with respect to hot-spot residues of native peptide ligands. *R*_*hot-spot*_ exceeds *R*_*all*_ by 6% approximately and equals to 51.07% on set T. The performance of the method is 1.5 times better on hot-spot residues of homo-oligomeric PPIs than of hetero-oligomeric PPIs: corresponding values of *R*_*hot-spot*_ are 57.75% and 37.19%. Recovery of hot-spot residues of subsets B-ab/ag and B-ag/ab is worse than on subset T-he. We got the lowest results on subset B-ag/ab, in large part due to poor recovery rates obtained on influenza’s hemagglutinin (16.95%, Table S5) which constitute 82% of complexes of this subset. At the same time, *R*_*hot-spot*_ observed on SARS-CoV-2 structures of subset B-ag/ab is equal to 100% (Table S5).

Assessing binding free energies 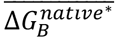 and 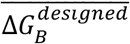 across test subsets, the binding sites have higher affinity for peptide ligands with native sequences binding energies on average, but the difference is small or equal to 0.7 REU on set T only (Table S6). We estimated complexes with native* peptides (see SI 1.2) instead of native ones during calculations of energetic metrics, for comparison with the designed complexes, in order to get rid of systematic superiority of native peptides due to more favorable backbone conformations which did not undergo perturbation. Comparison of native and designed by PepSeP1 method complexes in more details are discussed in sections SI 2.1-2.2. The accuracy of the predictions depends on relative solvent accessible surface area (SASA) of a given residue and Δ^i^G P-value of the native PPIs (Fig. S9). We observed some reduction of *R*_*all*_ with increasing of both SASA, as seen before (Chen et al., 2020), and Δ^i^G P-value. As expected, the decrease is more prominent for hetero-oligomeric PPIs. Results of application of PepSeP1 on concrete examples, including design of AAS for antibody fragments bound to SARS-CoV-2 and influenza’s HA are presented in SI 2.3.

As there can be different solutions to binding to the same interface (DeLano et al., 2000) we also integrated a variation of our software that produces six sequences, called PepSeP6. The output sequences differ on average within three positions usually. However, we also saw complete identical sequences or only 1 variation (Fig. S13). Such convergence is observed when all six outputs are generated with high rates of sequence recovery (80-100%).

*R*_*all*_ of the model across all six outputs is 39.54% (Table 3), which is lower than *R*_*all*_ of PepSeP1. However, *R*_*all*_ calculated only across the outputs most matching the native sequences out of six is equal to 52.8% (Table S9). The difference in binding energies of complexes with all six sequences is only within 1 REU in average (Fig. S14), the values are very close to native ones. Thus, PepSeP6 sequences provide comparable binding energies and could present alternatively binding solutions as seen in nature. More discussion of performance of PepSeP6 model is presented in SI 2.4.

The Rosetta FastDesign protocol is currently a commonly used protocols for the redesign of AAS demonstrating many experimentally validated designed protein interfaces (Cao et al., 2020; Huang et al., 2016; Jacobs et al., 2016; Linsky et al., 2020; Silva et al., 2019). Here, we compared the performance PepSeP1 model with FastDesign itself and with results of a combination these two methods in three versions, called RD3, RD5 and RD20, described in methods section. FastDesign results obtained using the default relax script are shown here (Table 3). Overall, redesigns of PepSeP1 sequences did not result in higher rates of recovery of native residues according to *R*_*all*_ values of RD3, RD5 and RD20 approaches. However, redesigns provide slightly higher rates *R*_*hot-spot*_ in comparison with PepSeP1 in case of hetero-oligomeric PPIs. PepSeP1 substantially outperformed FastDesign in sequence recovery when both are applied to the same all-glycine peptides. Rates of recovery obtained using InterfaceDesign2019 relax script are reported in Table S10.

HP peptide ligands having large RMSD values and indistinct subtle geometrical features which might have been used to identify residues (like dihedral angles characteristic for proline), but overall relevant for target binding sites nevertheless (Fig. S3), are very challenging targets for AAS design and recovery of hot-spot interactions comparatively to all-glycine peptides based on the native backbones. Thus, FastDesign incorporates an excessive number of glycine and proline residues to sequences for HP peptides (Fig. S16), the average binding energy is about −7 REU (Fig. S15) only, while 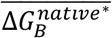 is −17.1 REU. Therefore, we developed PepSeP1 originally to work in conjunction with PepBB or on other peptide backbones engendered artificially for the *de novo* design of PPIs and having deviations from native backbones of different extents. The distribution of amino acid types in sequences predicted by PepSeP1 resembles the native distribution far better than FastDesign (Fig. S16), the average binding energy is −15.1 REU. Therefore, the performance of PepSeP1 on non-native peptide ligand fragments is far more relevant in comparison with FastDesign.

Figure 3 illustrate application of PepSeP6 method for design of AAS for HP peptide ligand generated for a highly conserved stalk region in HA and located with RMSD of 2.9 Å relative the CDR-loop of FI6 antibody; (the first design corresponds to PepSeP1 output). FI6 antibody fragment has three hot-spots at considered region: Arg99 (ΔΔ*G*_*i*_ = 3.2 REU), Leu100A (ΔΔ*G*_*i*_ = 4.1 REU), Tyr100C (ΔΔ*G*_*i*_ = 3.9 REU). Among PepSeP6 outputs, four designs contain recovered Leu100A interaction, two designs have Tyr near C-terminus and two designs have Arg near N-terminus; some of them provide two or more native-like recovered amino acids, as in complex selected for the illustration of the results. At the same time, FastDesign performed much poorer. Redesign of the complex obtained using PepSep6 method through RD20 approach allowed to keep native-like interactions involving Arg and Leu and to add recovered Ser residue.

**Figure 3.**
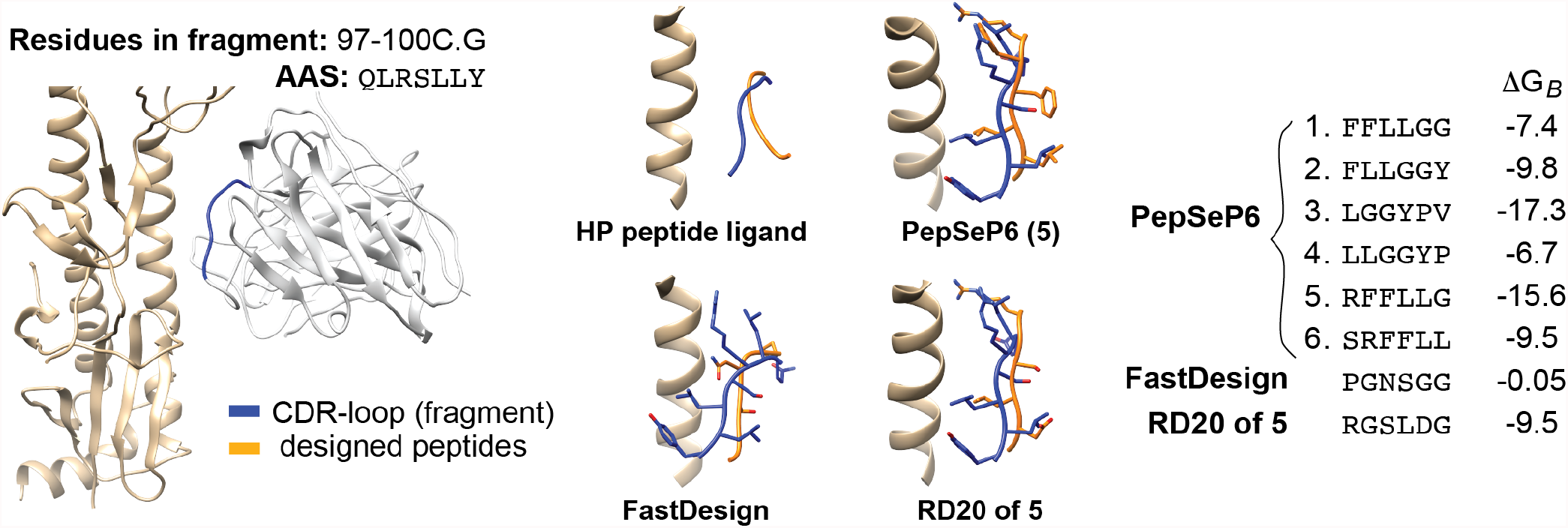
Design of highly perturbed (HP) peptide fragment generated as a binder for influenza’s hemagglutinin and located nearby CDR-loop of FI6V3 antibody structure (PDB code: 3ZTJ) by PepSeP6, FastDesign and RD20 methods. Δ*G*_*B*_ is in REU.

The results show that PepSep1 and PepSeP6 can reproduce relevant contacts; the resulted peptide ligands show high binding affinity and often outperform designs of FastDesign in case of redesigning a highly perturbed native backbones and thereby illustrates its usefulness for homology models as well as or potentially *de novo* designed backbones as they likely are not at the exact position they should be – as either methods have a high margin of error. Thus, a more knowledge-based design process can even guide the backbone refinement. The PepSeP6 method provides more diverse designs, iteration through which can reveal sequences with higher recovery rates, but it will be also useful for any affinity optimization processes as there are multiple solution to binding at a specific epitope.

## Conclusions

The neural network designed for recovery of peptide ligand sequences at a known protein binding site is performed in this study. To our knowledge, this is the first neural network model for prediction of amino acid sequences for peptides involved into interchain interactions. The model was developed in two versions: PepSep1 and PepSeP6 with single and multiple (six) outputs, correspondingly. The native sequence recovery rate of PepSep1 is 44.42% on the independent test set; the average accuracy of PepSeP6 designs is 39.54% with recovery rates of 52.8% across the output sequences resembling the native structures the most. The models characterized by intentional training on non-perfect backbones of structures which make them more applicable for non-perfect engineered *de novo* interactions, including homology models of PPI if one of the counterparts is intended for a redesign.

