## Supplementary material for "Deep learning of Protein Sequence Design of Protein-protein Interactions": SI

### 1 Methods

#### 1.1 Criteria for selection of PDB-files

PDB structures were selected from an entire PDB Bank in accordance with criteria:

1) structure does not contain DNA or RNA chains;

2) resolution is 2.0 Å or better if the crystal structure contains one protein entity or 2.5 Å or less if there are two or more entitles or 3.5 Å or less if it is an antibody-antigen complex;

3) the structure contains of 2 or more protein chains;

4) the structure is not an assembly of monomer molecules stabilized through crystal contacts only (the set of PDB files was analyzed using PISA protein-interface webserver(Krissinel & Henrick, 2007) and only those protein-protein interactions having $\Delta^i G$ P-value lower than 0.7 were selected);

5) molecular weight does not exceed 700 kDa;

6) ratio of the amino acids in α-helix structural motifs does not exceed 75%;

7) ratio of the amino acids in β-sheet structural motifs does not exceed 75%;

8) structures have 90% or less sequence identity.

Although $\Delta^i G$ P-value > 0.5 means that the interface is less hydrophobic than it could be, therefore the interface is likely to be an artefact of crystal packing, we found out that considerable amount of acknowledged antibody-antigen interfaces has P-value higher than 0.5. So, the threshold value of 0.7 was applied. The last three criteria were introduced to achieve the balanced dataset of ligand-binding site pairs, ideally without duplicates and with equal shares of loop, helix and sheet pieces.

#### 1.2 Preprocessing of input structures

PDB-files were preprocessed by means of clean_pdb.py script from Rosetta tools repository. The prepared protein structures were refined with Rosetta software by applying all-atom harmonic-restrained relax protocol with SD = 0.5.

For extraction of the ligand-binding site complexes from a crystal structure, a pool of potential ligand chains was determined first by randomly selecting single representatives per group of homologous chains. The ligand chains were cut to 6-residue fragments – ligands oligopeptides, with stride 1 residue (Fig. S1).

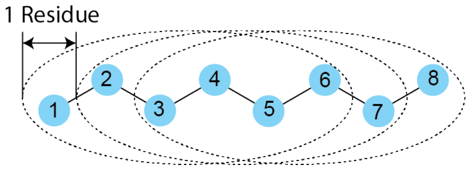


**Figure S1.** Generation of 6-residue peptide fragments from native protein ligands.

Binding pockets consists of 24-48 residues of the binding site chains which are closest to the ligand peptides when the distances between backbone atoms of peptide ligand residues and all atoms of surface residues are measured; the maximum distance threshold was set to 20 Å, the complexes having less than 24 residues within the threshold are not used. Average distances between peptides and binding sites of the complexes are presented in Fig. S2. Binding site chains are defined by types of the subsets in the case of the benchmark set and are all chains of the structure except the ligand chain in training, validation and test sets.

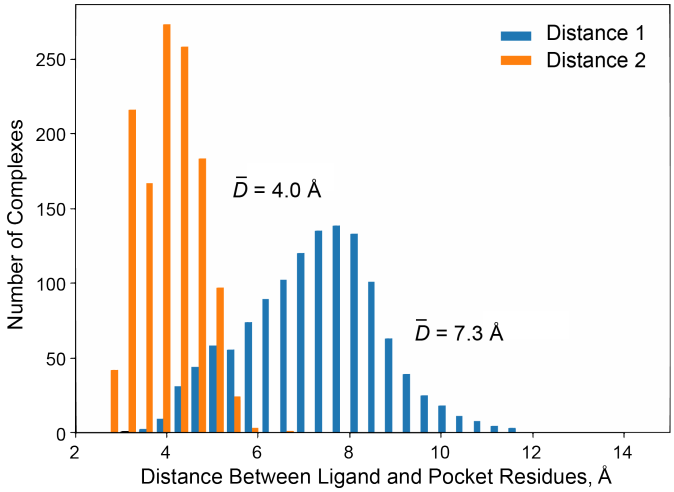


**Figure S2.** Ligand-pocket distances in complexes of the independent test set. Distance 1 – distance between coordinate center of Ca atoms of 4 central residues of peptide ligand and Ca atom of pocket residue closest to the coordinate center; distance 2 – average closest distance over backbone atoms of 4 central ligand residues till atoms of pocket residues; $\bar{D}$ – mean value.

Side-chain conformations of all residues were relaxed into the Rosetta force field using FastRelax protocol over 50 steps and subsequent MinMover over 100 steps. A few energetic features of the relaxed complexes are calculated: binding free energies $DG_{bind}$ are estimated by means of InterfaceAnalyzerMover, contributions $\Delta\Delta G_i$ of individual peptide residues $i$ to the binding are calculated by alanine scanning and pairwise interaction energies $\Delta\Delta G_{ij}$ between peptide residues $i$ and binding site residues $j$ are evaluated using scorefxn.eval_ci_2b function of Rosetta. Details can be found in methods section of SI.

The complexes for further consideration were selected according to following criteria thereafter:

1) two residues of a peptide ligand contribute to the binding with $\Delta\Delta G_i > 0.5$ REU and three nonterminal residues locate within 6 Å from a binding site when measuring distance between closest heavy atoms (side chains are taken into account) in the native interface at least, this distance threshold is denoted as *Thr;*
2) resolution thresholds for structures in subsets T-ho, T-he and the benchmark set are 2.0 Å, 2.5 Å and 3.5 Å, respectively.
3) non-standard residues and non-nutritional compounds should not be within 5 Å of fragments of interest;
4) the complexes should have negative binding free energies.

The current model did however not include complexes connected through covalent bonds or containing other atoms than found in the canonical amino acids.

Complexes were purged from duplicates before splitting into training, validation and test sets: if the ligands have the same amino acid sequences and the same six closest binding site residues located on comparable distances which vary within ± 1 Å, the peptide ligand complex with the lowest binding free energy was kept only. The benchmark set was purged separately in the same way. The complexes were reselected to make the minimal stride of two residues between the peptide ligands originating from the same chain in case of training and validation sets and three residues in case of test and benchmarking sets, based on the binding energies: the complex with the lowest binding energy was selected between two adjacent ones.

Distribution of complexes according to secondary structures of their peptide ligands in presented in Table S1.

**Table S1.** Fractions of oligomers with different types of secondary structure.

| Dataset | α-Helix | β-Sheet | Loop | Mixed |
|---|---|---|---|---|
| Training set | 32.06% | 17.86% | 24.85% | 25.23% |
| Validation set | 36.65% | 14.25% | 23.87% | 25.23% |

A peptide was assigned to a certain type if at least 4-residue sequence in it corresponds to that type (or, in the case of β-sheets, was corresponding in the original chain).

Before utilization, the peptide ligands were mutated to all-glycine binders and idealized by means of IdealizeMover. Furthermore, since, the model is intended to be used on engineered ligand backbones having some deviations from truly native poses because of problems with docking of bare backbones to the binding sites, the backbone conformations/poses were slightly distorted by small random movements within the binding sites using SmallMover over 25 steps, producing an average RMSD of 0.12 Å.

##### 1.3 Estimation of contributions of complex residues to binding

Alanine scanning performed for evaluation of contributions of individual amino acids of the peptide ligand was carried out without consideration of glycine, proline and alanine residues (substitutions of proline or glycine by alanine may cause a conformational change in the protein backbone and alanine to alanine substitution do not cause the energy change).

For some of our assessment metrics it is important to identify similar interactions in native and predicted complexes, but alanine scanning measures the total contribution of the residue to the binding free energy only without insights into formed interactions. Therefore, pairwise contributions of ligand and pocket residues were evaluated as a sum of two-body energies between the binding site residue and corresponding ligand residue by means of scorefxn.eval_ci_2b function of Rosetta using the following energy scoring terms:

$$DDG_{ij} = fa\_atr + fa\_rep + fa\_elec + fa\_sol \quad (S1),$$

where $DDG_{ij}$ is an interaction energy between the ligand residue *i* and the binding site residue *j*. Signs of $DDG_{ij}$ were changed to the opposite for more convenience in further applications.

**Table S2.** Benchmarking set of crystal structures

| No | PDB File | Title |
|---|---|---|
| | | *Coronavirus SARS-CoV-1* |
| 1 | 2dd8.pdb | Crystal structure of SARS-CoV spike receptor-binding domain complexed with neutralizing antibody |
| 2 | 3bgf.pdb | X-ray crystal structure of the SARS coronavirus spike receptor binding domain in complex with F26G19 Fab |
| 3 | 6waq.pdb | Crystal structure of the SARS-CoV-1 RBD bound by the cross-reactive single-domain antibody SARS VHH-72 |
| | | *Coronavirus SARS-CoV-2* |
| 1 | 6w41.pdb | Crystal structure of SARS-CoV-2 receptor binding domain in complex with human antibody CR3022 |
| 2 | 6yla.pdb | Crystal structure of the SARS-CoV-2 receptor binding domain in complex with CR3022 Fab |
| | | *Coronavirus MERS-CoV* |
| 1 | 4xak.pdb | Crystal structure of potent neutralizing antibody M336 in complex with MERS Co-V RBD |
| 2 | 4zpt.pdb | Structure of MERS-coronavirus spike receptor-binding domain (england1 strain) in complex with vaccine-elicited murine neutralizing antibody D12 (crystal form 1) |
| 3 | 5gmq.pdb | Structure of MERS-CoV RBD in complex with a fully human antibody MCA1 |
| 4 | 5yy5.pdb | Structural definition of a unique neutralization epitope on the receptor-binding domain of MERS-CoV spike glycoprotein |
| 5 | 6c6z.pdb | Crystal structure of potent neutralizing antibody CDC2-C2 in complex with MERS-CoV S1 RBD |
| 6 | 6nb3.pdb | MERS-CoV complex with human neutralizing LCA60 antibody Fab fragment (state 1) |
| | | *Influenza Virus Hemagglutinin* |
| 1 | 1eo8.pdb | Influenza virus hemagglutinin complexed with a neutralizing antibody |
| 2 | 1ken.pdb | Influenza virus hemagglutinin complexed with an antibody that prevents the hemagglutinin low pH fusogenic transition |
| 3 | 2vir.pdb | Influenza virus hemagglutinin complexed with a neutralizing antibody |
| 4 | 2vis.pdb | Influenza virus hemagglutinin, (escape) mutant with THR 131 replaced by ILE, complexed with a neutralizing antibody |
| 5 | 2vit.pdb | Influenza virus hemagglutinin, mutant with THR 155 replaced by ILE, complexed with a neutralizing antibody |
| 6 | 3fku.pdb | Crystal structure of influenza hemagglutinin (H5) in complex with a broadly neutralizing antibody F10 |
| 7 | 3gbm.pdb | Crystal structure of Fab CR6261 in complex with a H5N1 influenza virus hemagglutinin. |
| 8 | 3gbn.pdb | Crystal structure of Fab CR6261 in complex with the 1918 H1N1 influenza virus hemagglutinin |
| 9 | 3lzf.pdb | Crystal structure of Fab 2D1 in complex with the 1918 influenza virus hemagglutinin |
| 10 | 3ztj.pdb | Structure of influenza a neutralizing antibody selected from cultures of single human plasma cells in complex with human h3 influenza haemagglutinin. |
| 11 | 4fqi.pdb | Crystal structure of Fab CR9114 in complex with a H5N1 influenza virus hemagglutinin |
| 12 | 4fqj.pdb | Influenza B/Florida/4/2006 hemagglutinin Fab CR8071 complex |
| 13 | 4gms.pdb | Crystal structure of heterosubtypic Fab S139/1 in complex with influenza a H3 hemagglutinin |

| | | |
|---|---|---|
| 14 | 4hf5.pdb | Crystal structure of Fab 8F8 in complex a H2N2 influenza virus hemagglutinin |
| 15 | 4hfu.pdb | Crystal structure of Fab 8M2 in complex with a H2N2 influenza virus hemagglutinin |
| 16 | 4hg4.pdb | Crystal structure of Fab 2G1 in complex with a H2N2 influenza virus hemagglutinin |
| 17 | 4hkx.pdb | Influenza hemagglutinin in complex with CH67 Fab |
| 18 | 4hlz.pdb | Crystal structure of Fab C179 in complex with a H2N2 influenza virus hemagglutinin |
| 19 | 4lvh.pdb | Insight into highly conserved H1 subtype-specific epitopes in influenza virus hemagglutinin |
| 20 | 4m5z.pdb | Crystal structure of broadly neutralizing antibody 5J8 bound to 2009 pandemic influenza hemagglutinin, HA1 subunit |
| 21 | 4o58.pdb | Crystal structure of broadly neutralizing antibody f045-092 in complex with A/Victoria/3/1975 (H3N2) influenza hemagglutinin |
| 22 | 4py8.pdb | Crystal structure of Fab 3.1 in complex with the 1918 influenza virus hemagglutinin |
| 23 | 4r8w.pdb | Crystal structure of H7 hemagglutinin from A/Anhui/1/2013 in complex with a neutralizing antibody CT149 |
| 24 | 4ubd.pdb | Crystal structure of a neutralizing human monoclonal antibody with 1968 H3 HA |
| 25 | 4xnm.pdb | Antibody influenza H5 complex |
| 26 | 4xrc.pdb | Antibody hemagglutinin complexes |
| 27 | 4yk4.pdb | Human antibody 641 I-9 in complex with influenza hemagglutinin H1 Solomon Islands/03/2006 |
| 28 | 5a3i.pdb | Crystal structure of a complex formed between FLD194 Fab and transmissible mutant H5 haemagglutinin |
| 29 | 5dum.pdb | Crystal structure of influenza a virus H5 hemagglutinin globular head in complex with the Fab of antibody 65C6 |
| 30 | 5dup.pdb | Influenza a virus H5 hemagglutinin globular head in complex with antibody AVFLUIGG03 |
| 31 | 5dur.pdb | Influenza a virus H5 hemagglutinin globular head in complex with antibody 100F4 |
| 32 | 5gjs.pdb | Crystal structure of H1 hemagglutinin from A/California/04/2009 in complex with a neutralizing antibody 3E1 |
| 33 | 5gjt.pdb | Crystal structure of H1 hemagglutinin from A/Washington/05/2011 in complex with a neutralizing antibody 3E1 |
| 34 | 5ibl.pdb | Human antibody 6639 in complex with influenza hemagglutinin H1 X-181 |
| 35 | 5k9k.pdb | Crystal structure of multidonor HV6-1-class broadly neutralizing influenza A antibody 56.A.09 in complex with hemagglutinin Hong Kong 1968. |
| 36 | 5k9q.pdb | Crystal structure of multidonor HV1-18-class broadly neutralizing influenza A antibody 16.A.26 in complex with a/Hong Kong/1-4-MA21- 1/1968 (H3N2) hemagglutinin |
| 37 | 5kan.pdb | Crystal structure of multidonor HV1-18-class broadly neutralizing influenza A antibody 16.G.07 in complex with a/Hong Kong/1-4-MA21- 1/1968 (H3N2) hemagglutinin |
| 38 | 5ug0.pdb | Human antibody H2897 in complex with influenza hemagglutinin H1 Solomon Islands/03/2006 |
| 39 | 5ugy.pdb | Influenza hemagglutinin in complex with a neutralizing antibody |
| 40 | 5umn.pdb | Crystal structure of C05 VPGSGW mutant bound to H3 influenza hemagglutinin, HA1 subunit |

| | | |
|---|---|---|
| 41 | 5vag.pdb | Crystal structure of H7-specific antibody M826 in complex with the HA1 domain of hemagglutinin from H7N9 influenza virus |
| 42 | 5w08.pdb | A/Texas/50/2012(H3N2) influenza hemagglutinin in complex with K03.12 Fab |
| 43 | 5w0d.pdb | Inferred precursor (UCA) of the human antibody lineage K03.12 in complex with influenza hemagglutinin H1 Solomon Islands/03/2006 |
| 44 | 5w6g.pdb | Human antibody 6649 in complex with influenza hemagglutinin H1 Solomon Islands |
| 45 | 5wko.pdb | Crystal structure of antibody 27F3 recognizing the ha from A/California/04/2009 (H1N1) influenza virus |
| 46 | 5xhv.pdb | Crystal structure of Fab S40 in complex with influenza hemagglutinin, HA1 subunit. |
| 47 | 5y2l.pdb | Crystal structure of a group 2 HA binding antibody AF4H1K1 Fab in complex with the 1968 H3N2 pandemic (H3-AC/68) hemagglutinin |
| 48 | 6d0u.pdb | Crystal structure of C05 V110P/A117E mutant bound to H3 influenza hemagglutinin, HA1 subunit |
| 49 | 6e3h.pdb | Crystal structure of S9-3-37 bound to H5 influenza hemagglutinin |
| 50 | 6e56.pdb | Human antibody H2214 in complex with influenza hemagglutinin A/Aichi/2/1968 (X-31) (H3N2) |
| 51 | 6fyt.pdb | Structure of H1 (A/Solomon Islands/3/06) influenza hemagglutinin in complex with SD38 |
| 52 | 6fyu.pdb | Structure of H7(A/Shanghai/2/2013) influenza hemagglutinin in complex SD36 |
| 53 | 6fyw.pdb | Structure of B/Brisbane/60/2008 influenza hemagglutinin in complex with SD83 |
| 54 | 6ii8.pdb | Crystal structure of H7 hemagglutinin from A/Anhui/1/2013 in complex with a human neutralizing antibody l4B-18 |
| 55 | 6ml8.pdb | Crystal structure of hemagglutinin from H1N1 influenza a virus A/Denver/57 bound to the C05 antibody |
| 56 | 6mlm.pdb | H7 HA0 in complex with FV from h7.5 IGG |
| 57 | 6n5b.pdb | Broadly protective antibodies directed to a subdominant influenza hemagglutinin epitope |
| 58 | 6n5d.pdb | Broadly protective antibodies directed to a subdominant influenza hemagglutinin epitope |
| 59 | 6nz7.pdb | Crystal structure of broadly neutralizing influenza A antibody 429 B01 in complex with hemagglutinin Hong Kong 1968 |

**Table S3.** Quantities of complexes in subsets of benchmarking test sets according to their origin

| **Antigen** | **Subsets of Benchmarking Test Sets** | | | | |
|---|---|---|---|---|---|
| | B-ag/ag | B-ab/ab | B-ab/ag | B-ag/ab | **B** |
| SARS-CoV | 0 | 7 | 9 | 6 | 22 |
| SARS-CoV-2 | 0 | 8 | 6 | 1 | 15 |
| MERS-CoV | 0 | 40 | 15 | 19 | 74 |
| Hemagglutinin | 136 | 430 | 120 | 118 | 804 |
| **Total** | 136 | 485 | 150 | 144 | 915 |

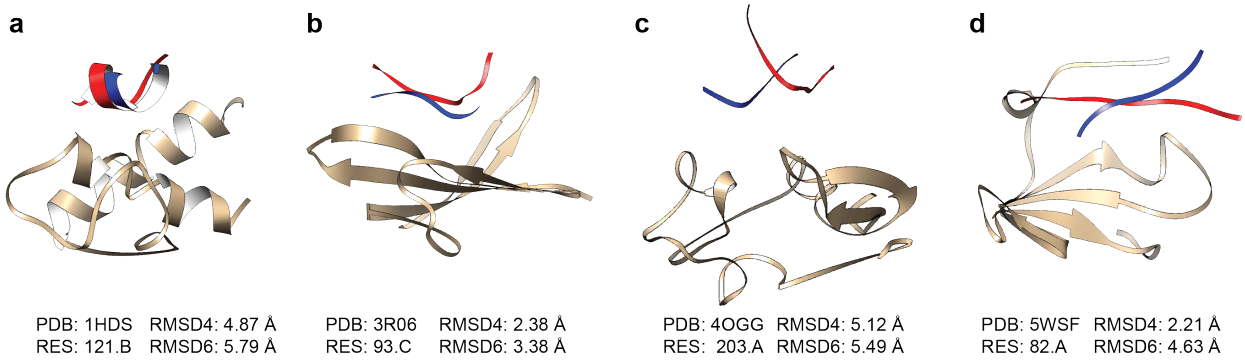


**Figure S3.** Visualization of random complexes from test set T with and native (red color) and designed by means of PepBB method (blue color) backbones of peptide ligands. Native structures are 8-mer fragments of protein ligands, designed structures consists of 6 residues. RES is a number of the second residue of the native protein ligand fragment in the original crystal structure. RMSD4 and RMSD6 are RMSD of the designed backbones taking into account non-terminal residues and all residues, respectively.

##### 1.4 Architecture of sub-models of the developed neural networks

The input for model 1 and model 2 was structured the following way:

1) Input 1 is an array of size 48 x 6 x 2 consisting of 2 distance maps of size 48 x 6. The distance maps consist of distances in Å between N and O backbone atoms of 48 binding site residues and 6 peptide ligand residues;
2) Input 2 is an array of size 48 containing secondary structure sequence of residues of the binding site.
3) Input 3 is an array of size 48 containing AAS of residues of the binding site.
4) Input 4 is an array of size 6 x 6 x 2 consisting of 2 distance maps of size 6 x 6 containing distances between backbone atoms of 6 residues of the peptide ligand; N and O backbone atoms are used for 2 distance maps, correspondingly.
5) Input 5 is a number (zero or one) labelling homo- or hetero-oligomeric type of PPI.
6) Input 6 is an array of size 6 which contains native AAS of the peptide ligand.
7) Inputs 7 is an analogue to input 1 but calculated for 20 residues of the binding site closest to the peptide ligands; size is 20 x 6 x 2.
8) Input 8 is an array of size 20 containing secondary structure sequence for 20 residues of the binding site used for input 7.
9) Input 9 of size 20 x 20 x 2 is an analogue to input 4 but calculated for 20 residues of the binding site used for input 7.

Zero padding is used if the number of the residues of the binding site is less than 48 in case of inputs 1-3.

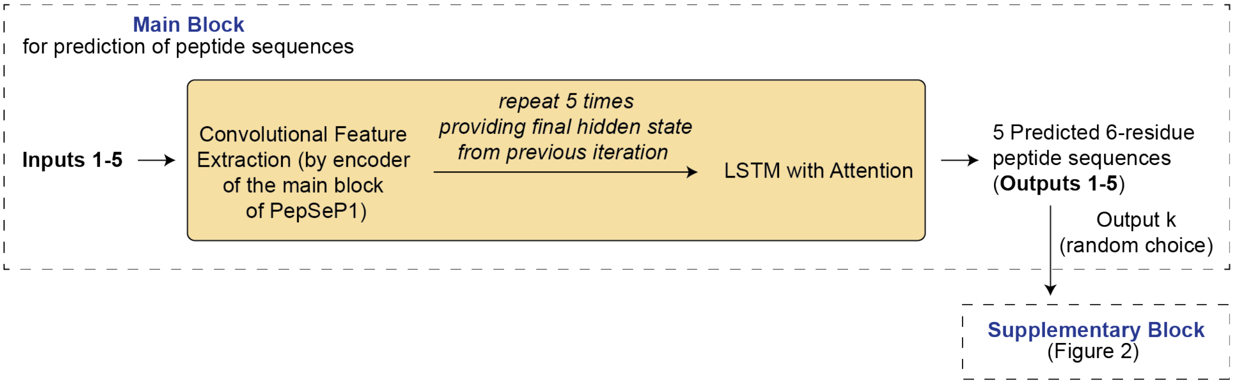


**Figure S4.** Scheme of PepSeP6 model, producing an ensemble of recovered amino acid sequences.

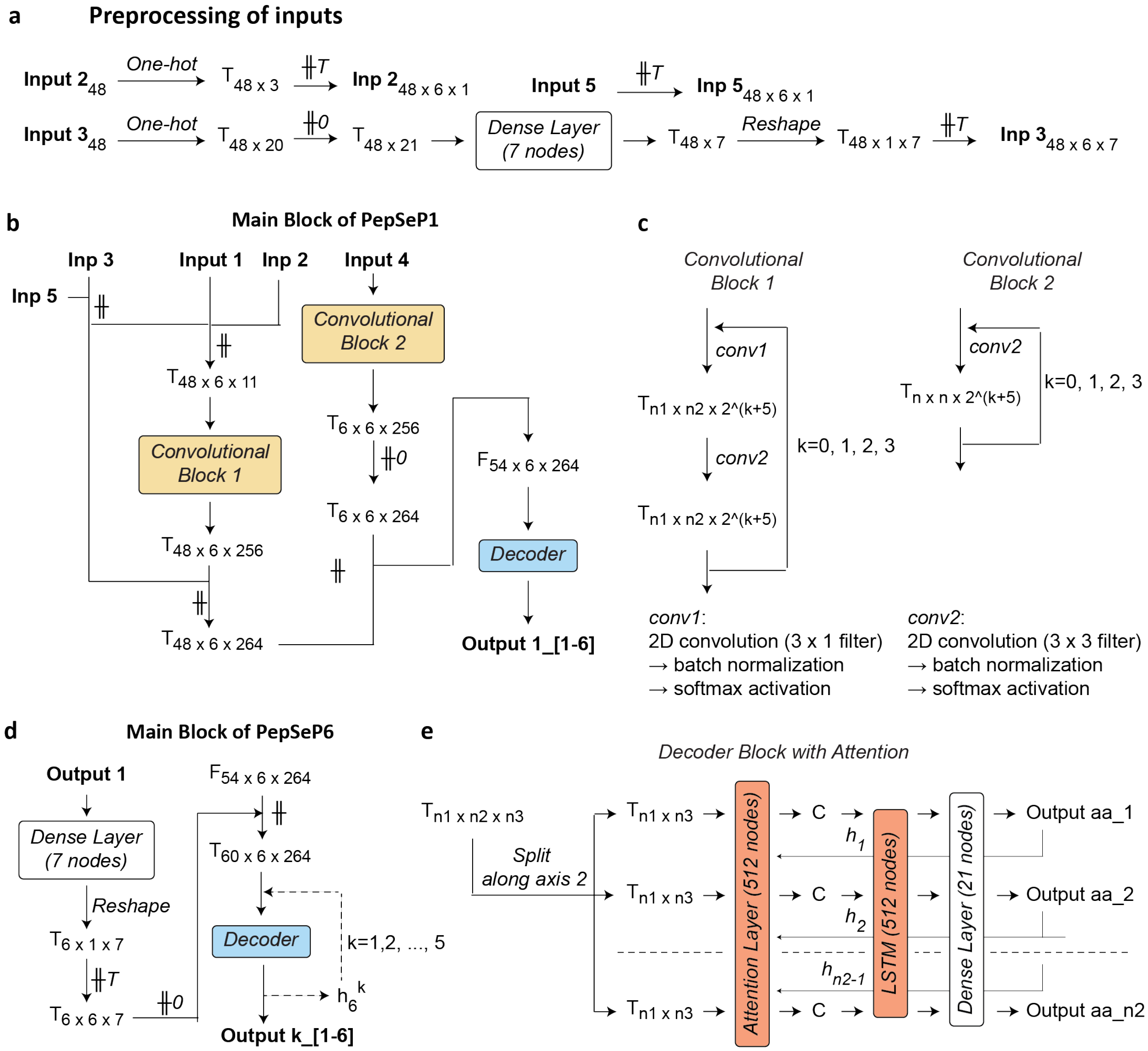

**Figure S5.** Architecture of the main blocks of PepSeP1 and PepSeP6 neural networks. (a) Preprocessing of inputs 2, 3 and 5. (b, d) General schemes of the main blocks of PepSep1 and PepSeP6, respectively; (c, e) architecture of convolutional blocks (c) and the decoder I. $T_{n1\ x\ n2\ x\ n3}$ is a tensor of size n1 x n2 x n3, $F_{54\ x\ 6\ x\ 263}$ is a tensor containing final feature vectors, C is context, ╫ is concatenate operation, ╫0 is zero-padding, ╫T is repeating of elements, output k_[1-6] is a recovered amino-acid sequence of 6-mer peptides with sequence number *k*, $h_i$ is a hidden state of LSTM layer after prediction of type of $i^{th}$ residue, $h_6^k$ is a hidden state of LSTM layer after prediction of the last residue of 6-mer peptide ligand with sequence number *k*, output aa_*i* is a predicted residue with sequence number *i*.

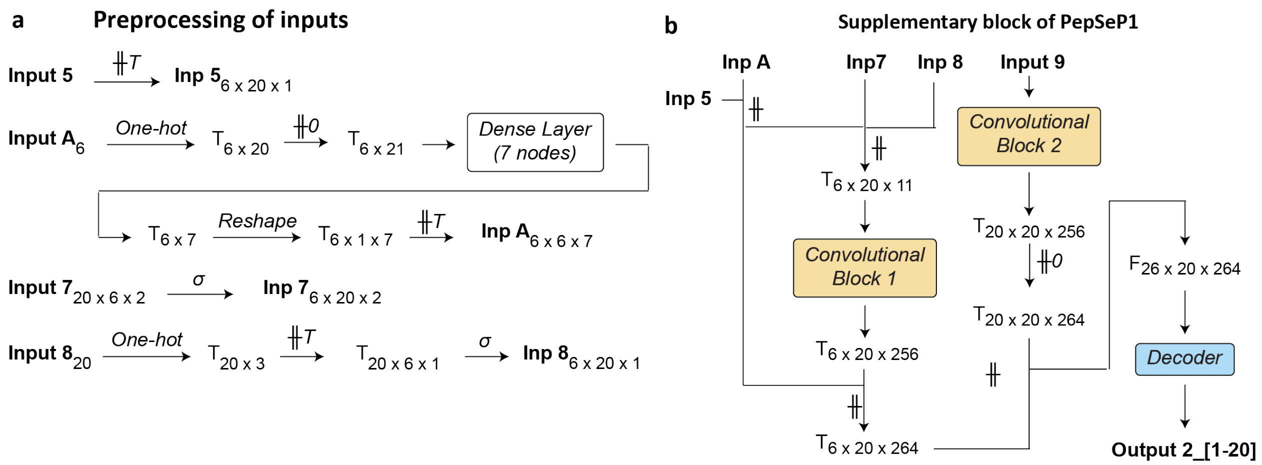


**Figure S6.** Scheme of the supplementary block of PepSeP1. (a) Preprocessing of inputs 5-8. (b) General scheme. Symbols corresponds to those of Fig.S3, σ is transpose operation; input A is amino acid sequence of the peptide ligand: native (input 6) or predicted (output 1).

Schemes of the main blocks of PepSep1 and PepSeP6 are presented in Fig. S5, the supplementary block of PepSeP1 is depicted in Fig. S6. Encoders utilize convolutional layers, there are two types of convolutional blocks were applied (Fig. S5c). Feature vectors extracted by both convolutional blocks are concatenated to form tensor F which second dimension equals to number of amino acids to predict. That tensor is an input of the decoder. Decoders of all blocks are similar attention-based recurrent neural networks, a general scheme of them is presented in Fig. S5e. Bahdanau-style (additive) attention with long-short-term memory (LSTM) units is used in the decoder. It follows the reference model (Image Captioning with Visual Attention, n.d.) with a few changes. The most notable difference with typical neural networks of such type is an absence of teacher forcing. This, due to relatively small number of canonical amino acids, does not affect the performance, but simplifies model and speeds up training. Sequential decoder inputs are types of amino acids predicted for preceding positions. Loss was calculated based on comparison of full target and predicted sequences. Encoder of PepSeP1 model was extracted and incorporated into PepSeP6 model without changes; during training the weights of the encoder were set as untrainable.

#### 1.5 Custom loss function of PepSeP6 model

Loss of the main block is a sum of average, maximum and minimum values of categorical cross entropy losses across all outputs of the ensemble, minimum value was scaled by 1.1. The scaling defines an amplitude between best and worst predictions in the ensemble: bigger values of scaling facilitate accuracy of the best predictions but the level of accuracy of the worst results is decreasing accordingly. An additional custom loss was introduced to the system to ensure diversity of the results, according formula:

$$custom\ loss = 1/\sum_{k1=1}^{5}\sum_{k2=1}^{5} CCE(o_{k1}, o_{k2})\ \ (S2),$$

where CCE is categorical cross entropy loss; $o_{k1}$, $o_{k2}$ are output sequences k1 and k2.

The custom loss calculates differences between outputs relative to each other using standard categorical cross entropy loss function; the final value is 1 divided by the sum of differences, that is, high diversity enhances smaller loss.

**1.6 Training of PepSep1 and PepSeP6 models**

Class- and sample- weighting was used during training. Class weights serve to compensate an unbalancing of the target classes, which in this work are types of canonical amino acids; the weights are inversely proportional to the frequency of the classes. Estimation of class weights was carried out using compute_class_weight function from scikit-learn library(Pedregosa et al., 2011). The resulting array was additionally raised to the power of 1.7 to make a difference between weights for rare and frequent amino acids more profound.

Sample weights intended to increase significance of amino acids with higher contribution into the binding energy during the network training. Computational $\Delta\Delta G_i$ results were obtained using alanine scanning for peptide ligands *i* and scorefxn.eval_ci_2b function for the residues of the binding sites. All residues with $\Delta\Delta G_i$ higher than 6 REU are weighted by 1. The weights for the residues having binding energy lower than 6 REU are calculated by normalization of the affinities to a range 0-1 according formula:

$$w = (x - min) / (max - min)\ \ (S3),$$

where w is a sample weight, x is $\Delta\Delta G_i$, *max* is 6, *min* is -2.

Residues with ΔΔG close to 6 REU are weighted by 1, and the opposite, the residues with low contributions are weighted by low values; *min* was set to -2 instead of 0 to make number of meaningful samples (with non-zero weights) more.

All convolution kernels were initialized using He's uniform initialization, LSTM layers utilized Glorot uniform initializer. Besides, both l1 and l2 regularization were applied in convolutional layers, the values were set to 1e-7 and 1e-6 correspondingly.

Training of PepSep1 model was carried out as follows:

1) Training of NN Sub-Model 2 utilizing amino acid sequences of native peptides (input A=input 6) ten times in three stages based on learning rates: 5 epochs with learning rate 0.001, 5 epochs with learning rate 0.0001 and 2 epochs with learning rate 0.00002. The trained model with the best performance was selected.

2) Training of NN Sub-Model 1 ten times through 5 epochs with learning rate 0.001. Three NN Sub-Models 1 with the best performance were selected.

3) Training of PepSep1 model with frozen weights of NN Sub-Model 2 which is fed by outputs of Sub-Model 1 (input A=output 1) and pretrained NN Sub-Model 1 through two stages : 5 epochs with learning rate 0.0001 and 2 epochs with learning rate 0.00001.

PepSeP6 model was trained 8 times using pretrained NN Sub-Models 1 and 2 in three stages based on learning rates: 5 epochs with learning rate 0.001, 5 epochs with learning rate 0.0001 and 2 epochs with learning rate 0.00002. PepSep1 and PepSeP6 models with best performances on the test sets were selected for future usage. Adam optimizer was used to minimize the mean squared error during optimization. Custom loss function was implemented for the training of PepSeP6, the details can be found in SI.

**1.7 Position-Specific Scoring Matrix (PSSM) scores calculation**

Scores for amino acid *aa* were calculated as following:

$$score(aa) = log_{10} \frac{P_{aa}^{NN}}{P_{aa}^{nat}} \text{ (S4)},$$

where $P_{aa}^{NN}$ and $P_{aa}^{nat}$ are probabilities that the residue at the position in question is of type *aa* according to the neural network model and the amino acid distribution in the native peptide ligands, respectively.

**1.8 Calculation of AI metric**

For estimation of AI, we first evaluated pairwise energies $DDG_{ij}$ between pocket residues and ligand residues of the native complex using scorefxn.eval_ci_2b, according equation S1. The pocket residues which total score

$$\Delta\Delta G_j = \sum_{i=1}^{6} \Delta\Delta G_{ij} \text{ (S5)},$$

is higher than 0.5 REU were considered further. The scores $DDG_{ij}$ for a given residue *j* were grouped and summed based on identity of the ligand residues, producing terms $\Delta\Delta G_{r-j}$, where *r* is an amino acid type, such as $\Delta\Delta G_{ALA-j}, \Delta\Delta G_{ARG-j}$, etc. The terms analogous for those were calculated for the designed complex as well. If at least a third of the native score $\Delta\Delta G_j$ is reproduced in the predicted complex due to the same $\Delta\Delta G_{r-j}$ terms, the pocket residue of the designed complex is counted as having the native bonds. In practice, for checking this condition, the following expression was used:

$$\sum_{r=1}^{20} |\Delta\Delta G_{r-j}^{n} - \Delta\Delta G_{r-j}^{d}| / \Delta\Delta G_j^{n} \leq 0.66 \text{ (S6)},$$

where superscripts *n* and *d* indicate native and designed complexes, respectively.

**1.9 Calculation of AS metric**

AS is calculated by comparing $\Delta\Delta G_j$ scores of the pocket residues: if both scores $\Delta\Delta G_j^n$ and $\Delta\Delta G_j^d$ are lower than 0.5 REU or the difference is less than 50% of the highest $\Delta\Delta G_j$ or $\Delta\Delta G_j^d$ is higher than $\Delta\Delta G_j^n$ , the residue is counted as a hit.

### 2 Supplementary Results

#### 2.1 Performance of PepSeP1 model

**Table S4.** Performance of PepSep1 method depending on an antigen type.

| Subset | Antigen | $R_{all}$, % | $R_{hot-spot}$, % |
|---|---|---|---|
| B-ab/ag | SARS-CoV-1 | 33.33 | 28.57 |
| | SARS-CoV-2 | 41.67 | 57.14 |
| | MERS-CoV | 32.22 | 31.25 |
| | HA | 26.53 | 32.89 |
| B-ag/ab | SARS-CoV-1 | 8.33 | 12.5 |
| | SARS-CoV-2 | 50.0 | 100.0 |
| | MERS-CoV | 26.32 | 35.0 |
| | HA | 16.95 | 21.84 |
| B-ag/ag | HA | 56.25 | 75.52 |
| B-ab/ab | SARS-CoV-1 | 71.43 | 60.0 |
| | SARS-CoV-2 | 87.5 | 100.0 |
| | MERS-CoV | 69.17 | 83.72 |
| | HA | 64.69 | 76.5 |

**Table S5.** Performance of PepSep1 method depending on secondary structure of the peptide ligands.

| Subset | Secondary structure | Number of samples | $R_{all}$, % | $R_{hot-spot}$, % |
|---|---|---|---|---|
| T-ho | α-helix | 337 | 53.71 | 61.14 |
| | β-sheet | 142 | 56.1 | 64.86 |
| | loop | 207 | 43.96 | 49.52 |
| | mixed | 184 | 52.63 | 54.02 |
| T-he | α-helix | 120 | 25.0 | 36.73 |
| | β-sheet | 77 | 30.52 | 40.7 |
| | loop | 95 | 29.3 | 37.27 |
| | mixed | 99 | 30.64 | 34.69 |
| T | α-helix | 457 | 46.17 | 54.41 |
| | β-sheet | 219 | 47.11 | 55.98 |
| | loop | 302 | 39.35 | 45.28 |
| | mixed | 283 | 44.94 | 47.06 |

A peptide ligand is assigned to a certain secondary structure if at least 4-residue sequence in it is of that type.

**Table S6.** Comparison of native and designed by PepSep1 method complexes.

| Subset | Number of samples | $\gamma_{interface}^{native^*}$, % | $\gamma_{interface}^{designed}$, % | $\gamma_{hot-spot}^{native^*}$, % | $\gamma_{hot-spot}^{designed}$, % | $\overline{\Delta G_B^{native^*}}$, REU | $\overline{\Delta G_B^{designed}}$, REU |
|---|---|---|---|---|---|---|---|

| | | | | | | | |
|---|---|---|---|---|---|---|---|
| *Test set* | | | | | | | |
| T-ho | 870 | 51.48 | 51.02 | 19.96 | 18.72 | -16.9 | -16.1 |
| T-he | 391 | 50.98 | 54.09 | 21.36 | 21.18 | -17.6 | -17.1 |
| **T** | **1261** | **51.32** | **51.97** | **20.39** | **19.48** | **-17.1** | **-16.4** |
| *Benchmark set* | | | | | | | |
| B-ab/ag | 150 | 46.22 | 42.56 | 21.22 | 16.67 | -16.0 | -13.1 |
| B-ag/ab | 144 | 45.95 | 49.31 | 16.44 | 16.67 | -13.8 | -13.4 |
| B-ag/ag | 136 | 56.5 | 55.51 | 27.08 | 28.19 | -16.7 | -16.2 |
| B-ab/ab | 485 | 44.64 | 46.77 | 22.82 | 23.51 | -15.3 | -15.5 |

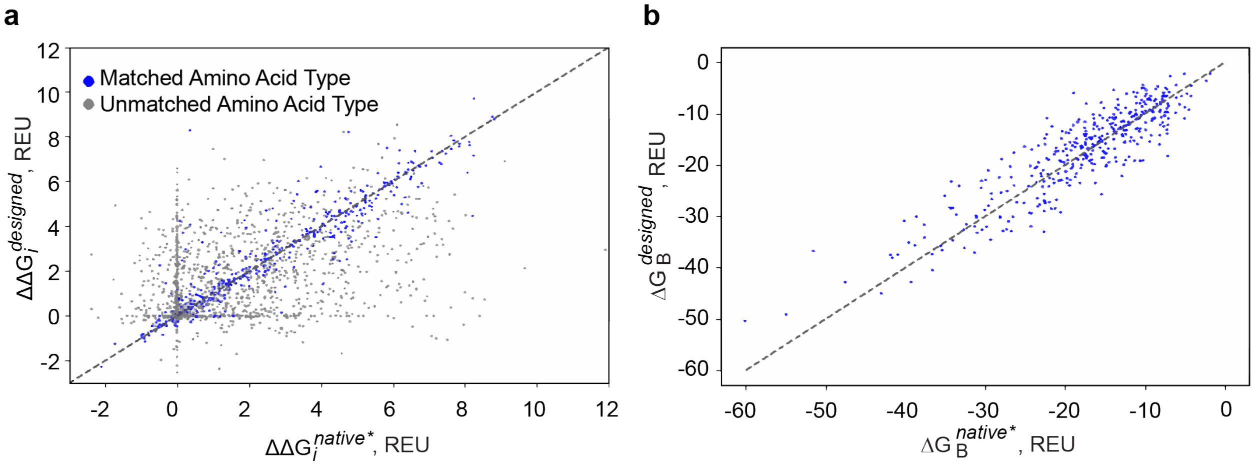


**Figure S7.** Correlation between energetic characteristics of complexes of test subset T-he (hetero-oligomeric PPIs) with native and designed by means of PepSeP1model peptide ligands: **(a)** binding free energies $\Delta\Delta G_i$ of residues with indication of matched identities, **(b)** binding free energies $\Delta G_B$ of complexes.

We defined interface residues alongside with hot-spot residues, the threshold of energy contribution to the binding for them was set to 0.3 REU which is 10 times less than for hot-spot residues (3 REU). We compared binding propensities of native and designed amino acid sequences by measuring the ratio of hot-spot residues $\gamma_{hot-spots}$ vs interface residues $\gamma_{interface}$ in complexed native* and designed peptides as quotement of number of interface or hot-spot residues to all residues of the peptides (Table S6, superscripts in names of metrics, e.g. $\gamma_{hot-spot}^{native*}$, stand for types of peptide ligand in the complexes) and by calculating binding free energies of the complexes $\Delta G_B$ by means of Rosetta's Interface Analyzer (Table S6, Fig. S7b). According to difference of 0.91% between $\gamma_{hot-spot}^{native*}$ and $\gamma_{hot-spot}^{designed}$ of set T, number of hot-spot residues in designed sequences slightly reduced relatively the native sequences. One the other hand, designed peptide sequences have slightly higher ratio of $\gamma_{interface}$ (0.65%) in case of set T. High contributions of the residues to the binding energy increase success of recapturing what is seen in natural interface, as it can be seen in Fig. S7a As it can be seen in Fig. 2b, the main range of the binding energies is comparable for both native and designed complexes. Overall closeness of the binding affinities indicates a reasonable quality of the predicted sequences, fitted both to a given secondary structure of the peptide and the binding surface.

Binding site residues in a designed complex can be involved in interactions with residues of designed peptides using the same kind interactions (e.g. salt bridges or van der Waals), but not with the residue at the original position: for example, ARG of the binding site is interacting with ASP at the first position of the peptide in the native complex but with GLU predicted for third position in the designed complex. Therefore, we analyzed similarity of the formed interactions in the complexes by means of IMI (identity matching interactions) and FMI (functionally matching interactions) metrics (Table S7). Both of them is a percentage of the ratio between number of the binding site residues of the designed complexes involved into interactions similar to those which are observed for them in the native complexes to the total number of interface binding site residues counted in the native complexes, the interacting binding site residues are selected using 0.3 REU threshold for DDG; IMI takes into account the interactions with exactly matched amino acid identities of the peptide ligands and FMI counts residues having side chains similar to the native ones as well (e.g. ASP and GLU; a classification of canonical amino acids based on their structure similarity is provided in Table S8). Additional metric EMI (energetically matching interactions) is a share of the binding site residies involved into interactions which energetically better or comparable with original scores but with ligand residues which do not resemble native types at all. More details about metrics can be found in S-1.8 and S-1.9.

**Table S7.** Recovery rates of native-like interactions achieved by PepSep1 method.

| Subsets | Interacting residues, % | | | Hot-spot pocket residues, % | | |
|---|---|---|---|---|---|---|
| | *Exact match* (IMI) | *Chemically similar match* (FMI) | EMI | *Exact match* (IMI) | *Chemically similar match* (FMI) | EMI |
| T-ho | 58.58 | 63.61 | 18.02 | 66.5 | 73.74 | 18.21 |
| T-he | 38.12 | 49.95 | 25.67 | 50.59 | 64.17 | 26.49 |
| **T** | **52.04** | **59.25** | **20.47** | **61.35** | **70.64** | **20.89** |
| B-ab/ag | 29.19 | 40.18 | 25.05 | 31.14 | 47.91 | 31.14 |
| B-ag/ab | 22.38 | 38.14 | 30.69 | 32.12 | 55.76 | 25.45 |
| B-ag/ag | 64.58 | 70.55 | 13.33 | 72.68 | 80.88 | 8.74 |
| B-ab/ab | 72.23 | 79.0 | 9.14 | 76.69 | 87.24 | 8.98 |

**Table S8.** Division of canonical amino acids into groups depending on their properties.

| **Group** | **Amino Acid Type** | **Group** | **Amino Acid Type** | **Group** | **Amino Acid Type** | **Group** | **Amino Acid Type** |
|---|---|---|---|---|---|---|---|
| 1 | ASP, GLU | 4 | GLY | 7 | PRO | 10 | TYR, PHE, TRP, HIS |
| 2 | ASN, GLN | 5 | ALA, CYS | 8 | VAL | 11 | HIS, ARG |
| 3 | ILE, LEU | 6 | MET | 9 | SER, THR, TYR | | |

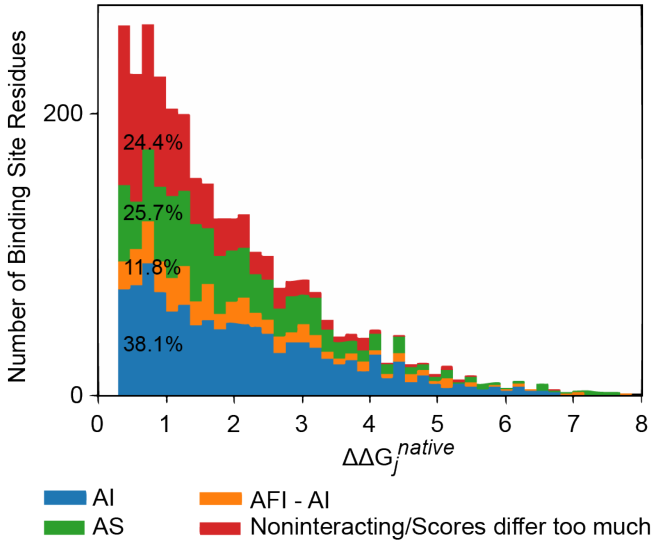


**Figure S8.** Dependence of rates of recovery of native interactions in the designed complexes of subset T-he on contribution of the binding site residues to the binding energy.

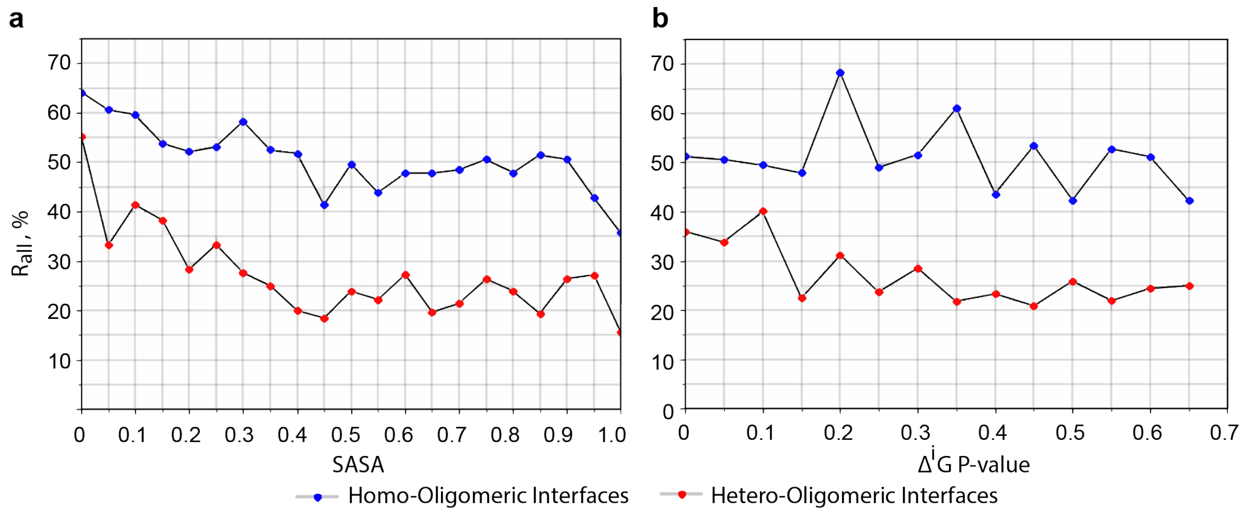


**Figure S9.** Rates of recovery $R_{all}$ for residues depending on relative sidechain SASA (**a**) and P-values of PPIs (**b**) on samples of set T.

Values of IMI exceed $R_{all}$ by 7.62% in set T, but it is due to accounting of interacting pocket residues only mostly whereas $R_{all}$ takes into account recovery of both interface and non-interfaces residues of peptide ligands. The binding site residues displaying higher contributions to the binding in the native structures are found to be involved into the same interactions in the designed complexes more frequently (Fig. S8). Counting of interactions based on similarity of side chains of ligand residues through FMI metric provides higher rates of the recovered interactions, as expected. The difference between IMI and FMI is bigger in case of hetero-oligomeric PPIs: values of FMI on interacting and hot-spot residues are 49.95% and 64.17% on subset T-he, respectively, which exceeds values of IMI by 10%. The difference on subsets B-ab/ag and B-ag/ab is even higher, and it is interesting that the performance of the method on hot-spot residues of subset B-ag/ab is better than on subset B-ab/ag. Considering metric EMI, it is demonstrated that 65-90% of interacting binding site residues are involved into native-like interactions or interactions energetically close to those of the native complexes.

##### 2.2 Estimation of selectivity of binders

16 Subsets of complexes consisting of six complexes with different ligands but with similar binding sites were prepared: 8 subsets from test subset $t_{ho}$ and 8 subsets from test subset $t_{he}$. Complexes within a subset should comply with the following criteria: four binding site residues out of six, closest to the peptide ligand, have to have matching amino acid types and the pairwise distances between these residues and ligand backbone atoms should differ no more than 2.0 Å in average.

The ligand sequences of the subset were tested against each of the subset pockets by mutation of original residues to designs obtained by PepSP1 and FastDesign protocol and subsequent relaxation using FastRelax according to usual routine described in methods section of the main paper. The measured binding affinities were ranked from 1 (best) to 6 (poorest) and the average positions in the ranking for the designs are reported. The summarized results are given in Fig. S10.

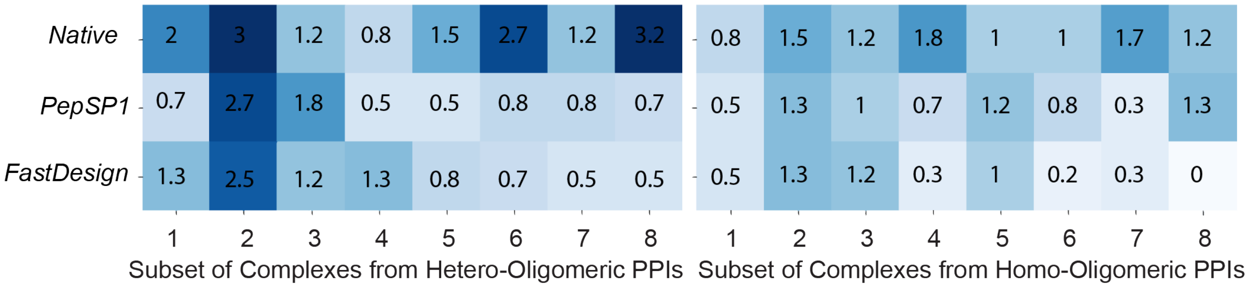


**Figure S10.** Average ranking of the binding energies of the complexes between the binding energies obtained while mutating the peptide ligand sequence to the sequences of the peptide ligands of other five complexes with similar but not identical binding sites

The average ranks for all types of sequences are close and adequately high in most of the cases. The best results are obtained on the native and FD sequences, PepSP1 sequences appeared to be less selective, although the rankings do not differ too much, especially for the hetero-oligomeric PPIs. Based on these results, we can summarize that the selectivity of the designed sequences is acceptable and in some cases is better than those of the native structures.

#### 2.3 Case study: Antibody – antigen interactions

The antibody CR3022 binds to the receptor binding domain (RBD) of the Severe Acute Respiratory corona virus 2 (SARS-CoV-2) (Fig. S11). To evaluate sequence recovery, we extracted each loop of the heavy chain that is forming contacts with the RBD in form of 6-residues fragments. After perturbation of these contacts and changing them to poly-glycine residues, we redesigned each fragment using PepSeP1. For redesign we saw that $R_{all}$ was more or equal to 50% in case of first two designs obtained by means of PepSeP1; interestingly, the Arg that was introduced at position 30 fits appears to be a better contact as it points into a highly negatively charged area, including the C-terminus of a helix (residues 365 – 371, Fig. S12). For the CDR3, most contacts are not contributing to much of the binding energy and accordingly, we only see an $R_{all}$ of about 16%. However, residues at 3$^{rd}$ and 4$^{th}$ positions in both native and designed amino acid sequences have functional similarity. PepSeP1 outperformed FastDesign in all three cases regarding $R_{all}$ as well as $\Delta G_B$ values.

The 5J8 antibody is a broadly neutralizing antibody that binds to residues within the receptor binding site of influenzas hemagglutinin (HA). Its main contact CDR-loop exceeds 6 residues, so, we performed PepSeP1 design by aligning two designs predicted for 6-residue fragments. The antibody has two residues strongly contributing to the binding with HA1 subunit over positions 97-100E of chain H: Tyr100 ($\Delta\Delta G_i$ = 3.7 REU) and Asp100B ($\Delta\Delta G_i$ = 8.2 REU). We can see that PepSep1 successfully recovers Asp at position 100B which mimics the carboxy group of sialic acid and the design contains aromatic TRP at position 100 which establishes a large contact area with the aliphatic part of Lys133 of the HA molecule. Besides these, we saw recovery of Ser98 and Pro100A. well. FastDesign recovered the same residues, however, Tyr or another aromatic or hydrophobic residue at position 100 is not predicted.

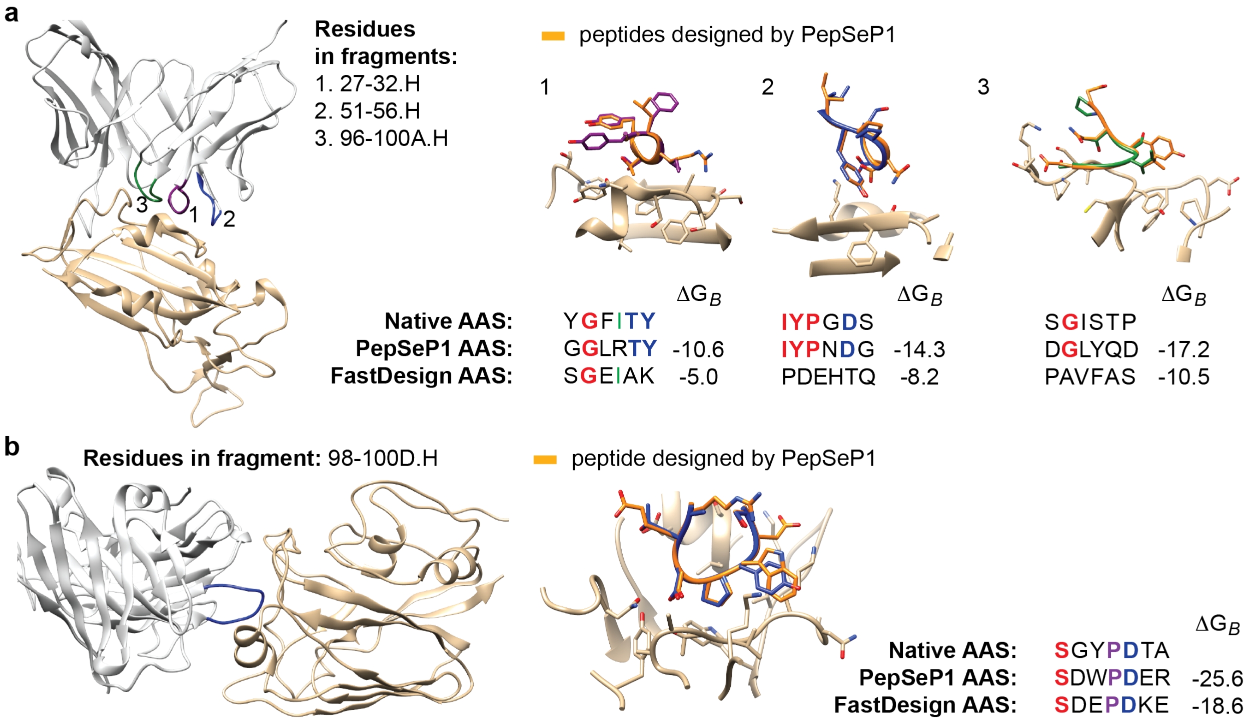


**Figure S11.** Native and designed fragments of antibody-antigen interfaces and corresponding amino acid sequences (AAS). (**a-b**) Design of fragments of CR3022 antibody complexed with

SARS-CoV-2 (PDB code: 6W41, **a**) and of 5J8 antibody complexed with influenza's hemagglutinin (PDB code: 4M5Z, **b**) by PepSeP1 and FastDesign methods.

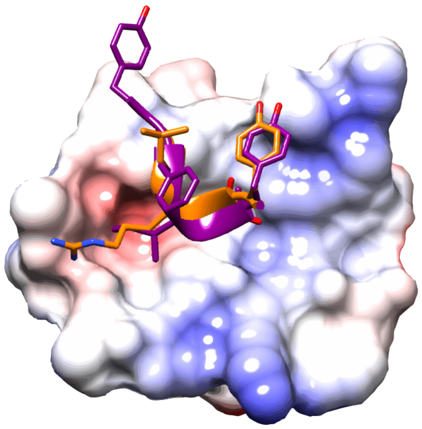

**Figure S12.** Native (purple) and designed (orange) fragments of CR3022 antibody (Fig. S11a, 1) complexed with SARS-CoV-2 (PDB code: 6W41) which surface is colored according to electrostatic potential.

##### 2.4 Performance of the PepSeP6 model

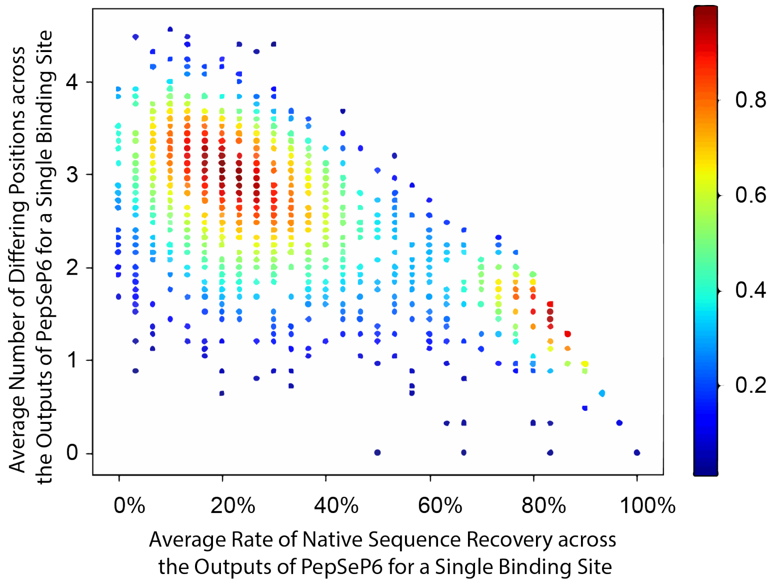


**Figure S13.** Dependence of diversity of outputs of PepSeP6 model on accuracy of the outputs with showing density distribution.

PepSeP6 model produces six sequences for the peptide ligands. The sequences resembling the native ones in the most and the least extent are marked as $O_1$ and $O_6$, respectively, and the sequences displaying the highest binding affinity are denoted as $O_e$.

**Table S9.** Native sequence recovery rates achieved by PepSeP6 on subsets of test and benchmarking sets in case of $O_1$, $O_6$ and $O_e$ sequences

| Subset | $O_1$ sequences | | $O_6$ sequences | | $O_e$ sequences | |
|---|---|---|---|---|---|---|
| | $R_{all}$, % | Rh, % | $R_{all}$, % | Rh, % | $R_{all}$, % | Rh, % |
| *Test set* | | | | | | |
| T-ho | 58.91 | 68.04 | 20.44 | 23.22 | 42.07 | 54.89 |
| T-he | 39.22 | 52.89 | 15.05 | 15.17 | 28.77 | 39.92 |
| **T** | 52.8 | 63.12 | 18.77 | 20.61 | 37.95 | 50.03 |
| *Benchmark set* | | | | | | |
| B-ab/ag | 38.44 | 52.36 | 13.78 | 15.18 | 25.56 | 35.6 |
| B-ag/ab | 28.7 | 37.32 | 6.94 | 9.15 | 15.39 | 23.94 |
| B-ag/ag | 63.48 | 80.09 | 19.12 | 26.24 | 46.08 | 66.06 |
| B-ab/ab | 71.34 | 82.23 | 52.06 | 65.21 | 60.69 | 76.66 |

Recovery rates for the sequences are presented in Table S9. Accuracy $R_{all}$ on the complexes of set T in case of $O_1$ sequences is 52.8%; the rates exceed the results of PepSeP1 method by 7.236% and 10.66% for homo-oligomeric and hetero-oligomeric PPIs, respectively. The differences are higher in case of hot-spot residues, especially for hetero-oligomeric PPIs: improvement for complexes of subset T-he is 15.7%, and the results are higher by 18.84% for antigen-antibody interfaces of subset B-ab/ag.

Recovery rates for $O_6$ and $O_e$ sequences of set T are 18.77% and 37.95%, correspondingly. Although the accuracies of $O_6$ sequences are lower than the results obtained on $O_1$ sequences by 20-30%, the difference in the binding energies of $O_1$ and $O_6$ sequences is within 1 REU only (Fig. S14). Thus, despite the low rate of hits, $O_6$ sequences provide the interactions with binding energies comparable with those of $O_1$ sequences. Both types of sequences demonstrate less reactivity than the native structures, although the deviation is small (0.2-1.0 REU). Sequence recovery rates for $O_e$ sequences (37.95%) are below average values across all six outputs on most subsets (39.54%, Table 2), although hot-spots recovery rates are superior and equal to 50.03% on set T. However, the highest $R_{hot-spot}$ is obtained on $O_1$ sequences still. Therefore, during the design of the peptide ligands by mean of PepSeP6, it is reasonable to consider all sequences and do not confine to the most energetically favorable only.

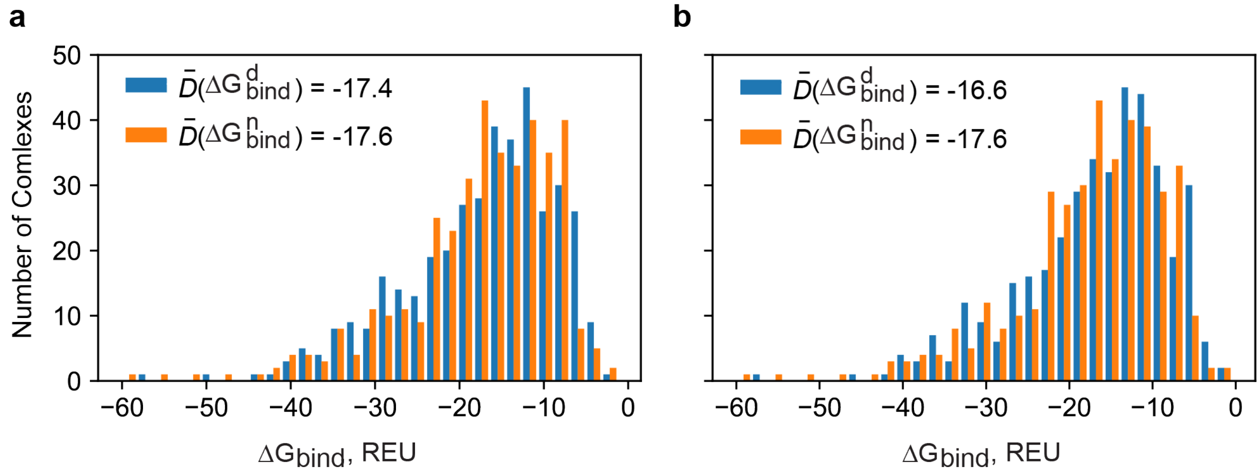

**Figure S14.** Distributions of the binding energies of complexes of test subset T-he with native and designed by PepSeP6 model peptide ligands: (a) designed peptide sequence is $O_1$ sequence, (b) designed peptide sequence is $O_6$ sequence.

#### 2.5 Comparison with performance of Rosetta's FastDesign protocol

**Table S10.** Rates of recovery of native residues of peptide ligands of sets T and B obtained by means of RD3, RD5, RD20 and FastDesign approaches, where InterfaceDesign2019 relax script utilized with FastDesign protocol

| Subset | $R_{all}$, % | | | | $R_{hot\text{-}spot}$, % | | | |
|---|---|---|---|---|---|---|---|---|
| | RD3 | RD5 | RD20 | FastDesign | RD3 | RD5 | RD20 | FastDesign |
| T-ho | 42.2 | 37.61 | 30.61 | 23.3 | 56.11 | 56 | 51.75 | 44.1 |
| T-he | 28.13 | 27.15 | 24.47 | 25.06 | 42.86 | 44.67 | 42.4 | 45.58 |
| **T** | 37.84 | 34.36 | 28.71 | 23.84 | 51.81 | 52.32 | 48.71 | 44.58 |
| B-ab/ag | 27.44 | 27.22 | 23.22 | 23.22 | 39.01 | 38.46 | 35.71 | 32.97 |
| B-ag/ab | 18.75 | 21.41 | 18.06 | 23.96 | 25.86 | 32.76 | 30.17 | 54.31 |
| B-ag/ag | 44.73 | 37.99 | 31.13 | 20.22 | 69.79 | 65.62 | 59.9 | 30.73 |
| B-ab/ab | 53.47 | 46.46 | 37.94 | 24.54 | 73.7 | 72.15 | 66.78 | 48.96 |

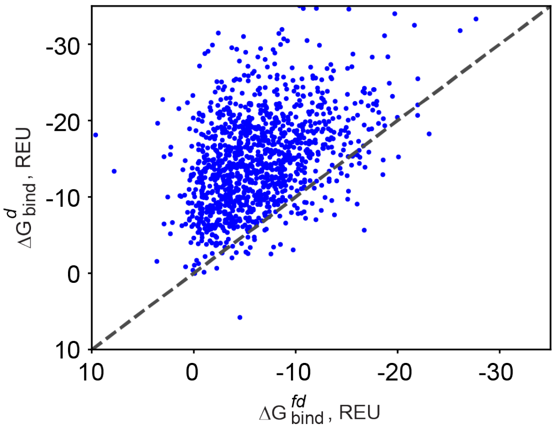


**Figure S15.** Binding free energies of complexes with highly disturbed peptide ligands (RMSD of non-terminal residues is 3.9 Å) of test set T, which amino acid sequences are designed by PepSep1 ($\Delta G^{d}_{bind}$) and FastDesign ($\Delta G^{fd}_{bind}$) methods.

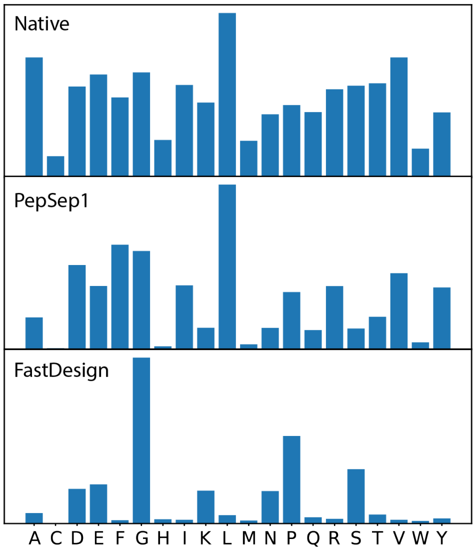


**Figure 16.** Amino acid distributions in HP peptide ligands with native and designed by PepSeP1 and FastDesign methods amino acid sequences.

Metrics IMI, FMI and EMI measured on hot-spot binding site residues of PepSeP1 designed complexes with HP peptides of subset T-he are 17.5%, 30.0% and 22.7%, respectively; these metrics were not measured in case of FastDesign due to its very poor performance.
